# Transiently Worse Postural Effects After Vestibulo-ocular Reflex Gain-Down Adaptation in Healthy Adults

**DOI:** 10.1101/2024.04.09.588711

**Authors:** Cesar Arduino, Michael C Schubert, Eric Anson

## Abstract

Suffering an acute asymmetry in vestibular function (i.e. vestibular neuritis) causes increased sway. Non-causal studies report associations between lateral semicircular canal function and balance ability, but direct links remains controversial. We investigate the immediate effect on body sway after unilateral vestibulo-ocular reflex (VOR) gain down adaptation simulating acute peripheral vestibular hypofunction. Eighteen healthy adults, mean age 27.4 (± 12.4), stood wearing an inertial measurement device with their eyes closed on foam before and after incremental VOR gain down adaptation to simulate mild unilateral vestibular neuritis. Active head impulse VOR gain was measured before and after the adaptation to ensure VOR gain adaptation. Percentage change for VOR gain and sway area were determined. Sway area was compared before and after VOR adaptation. VOR gain decreased unilaterally exceeding meaningful change values. Sway area was significantly greater immediately after VOR gain down adaptation, but quickly returned to baseline. In a subset of subjects VOR gain was re-assessed and found to remain adapted despite sway normalization. These results indicate that oculomotor adaptation targeting the lateral semicircular canal VOR pathways have an immediate, albeit transient increase in body sway. Rapid return of body sway to baseline levels suggests dynamic sensory reweighting between vestibular and somatosensory inputs to resolve the undesirable increased body sway.

## Introduction

Vestibular inputs have long been recognized as contributing to postural control (Nashner 1971; Lacour et al. 2022), with some studies implicating the otoliths as the primary vestibular contributors to postural control (Markham 1987; Diaz-Artiles and Karmali 2021). Galvanic vestibular stimulation causes postural tilts and acts by stimulating the vestibular nerve, likely presenting a combined canal-otolith signal to postural pathways (Fitzpatrick and Day 2004; Cathers et al. 2005). Other studies have correlated measures of lateral semicircular canal VOR function with body sway, especially using an eyes closed on foam test to increase the postural demands on the vestibular system (Anson et al. 2017, 2019). A recent paper described a model that accounts for separate signaling from semicircular canals (SCC) and otoliths to characterize postural control (Haggerty et al. 2017), with the SCC effects described as detecting transients. Roll-tilt perception involving the otoliths and vertical SCC has been associated with postural control, but horizontal SCC perception was not (Karmali et al. 2017; Beylergil et al. 2019). Interestingly, neither roll plane nor pitch plane VOR gain asymmetries were associated with balance, but rather roll VOR gain asymmetries were associated with gait abnormalities (Allum and Honegger 2020). VOR gain asymmetries associated with gait are in line with the suggestion that the SCCs role in postural control are detecting transients, i.e. heel strike, but this would argue against a prominent role in quiet stance postural control. It remains controversial whether there is a direct link between lateral SCC function and postural control.

Technology advancements in gaze stabilization exercises, specifically incremental VOR adaptation, offer both a chance at recovery of VOR function for patients (Gimmon et al. 2019; Rinaudo et al. 2021), and also a mechanism to probe motor learning within vestibular pathways (Gimmon et al. 2018). VOR adaptation is maximized following a 15 minute incremental VOR adaptation protocol (Muntaseer Mahfuz et al. 2018), and has been shown to last for at least to 60 minutes (Mahfuz et al. 2018). To date, the emphasis of studies examining incremental VOR adaptation has been restricted to the VOR pathway. In a single clinical trial for individuals with unilateral vestibular hypofunction walking balance improved after one week (Rinaudo et al. 2021), again suggesting semicircular canal function is important for walking. Standing balance was not assessed, and it is not known whether VOR adaptation will have any immediate effect on balance ability.

In healthy adults incremental VOR results in adaptation around 10-20% (Gimmon et al. 2018; Schubert and Migliaccio 2019), approaching the threshold of VOR decline noted in presbyvestibulopathy (Agrawal et al. 2019). In this study, we examined body sway with eyes closed on foam before and after incremental VOR gain down adaptation as a model to simulate mild unilateral semicircular canal only hypofunction on postural control in healthy subjects. We hypothesized that after incremental VOR gain down adaptation body sway would increase.

## Methods

### Subjects

Eighteen healthy adults (12 females, 6 males) mean age 27.4 (± 12.4), provided written informed consent and participated in this study, approved by the Institutional Review Board at the University of Rochester Medical Center. Participants were compensated $15 per hour.

### Experimental Setup

#### Equipment

VOR gain was measured using an EyeSeeCam head impulse system (Micromedical, Inc.) (Bartl et al. 2009; Schneider et al. 2009), using active head impulses in the plane of the horizontal SCC performed while seated in the dark. A projected world fixed laser dot on the wall that extinguished during head movement served as the fixation point. StableEyes ® was used to generate both the stable fixation point used for pre- and post-head impulse testing and provided the incremental retinal slip error signal during the 15-minute adaptation period (Muntaseer Mahfuz et al. 2018). Body sway was measured using an OPAL sensor (APDM, Inc) (Mancini et al. 2012).

#### Protocol

After consenting, each participant provided demographic data including age, gender, and fall history. Participants completed 2-3 training sessions at least a week before the experimental session to learn the correct technique for performing active head impulses (peak head velocity between 120 and 300 degrees/sec within 100ms) and a separate experimental session verifying that they were successful at VOR adaptation using the incremental VOR adaptation protocol. Visual and auditory feedback were provided to the subjects during the training sessions using the EyeSeeCam (visual feedback) and StableEyes ® (auditory feedback) systems.

After verifying that subjects could adapt their VOR with active head impulses, they were invited to participate in this experiment. During this experiment, subjects wore an OPAL sensor (APDM, Inc) on a belt around their waist. Subjects also wore both an EyeSeeCam system and StableEyes ® system on their head. Balance was measured with subject’s eyes closed standing on a foam balance pad (AIREX, Inc. density 38.6 kg/m^3) with their arms crossed and feet angled 15 degrees out from midline and separated approximately 10cm at the heels according to the APDM manual. Subjects completed three 30-second trials standing in the dark with eyes closed on the foam both before and after completing seated incremental VOR adaptation, see Figure 1 for experimental schematic.

**Figure 1.**
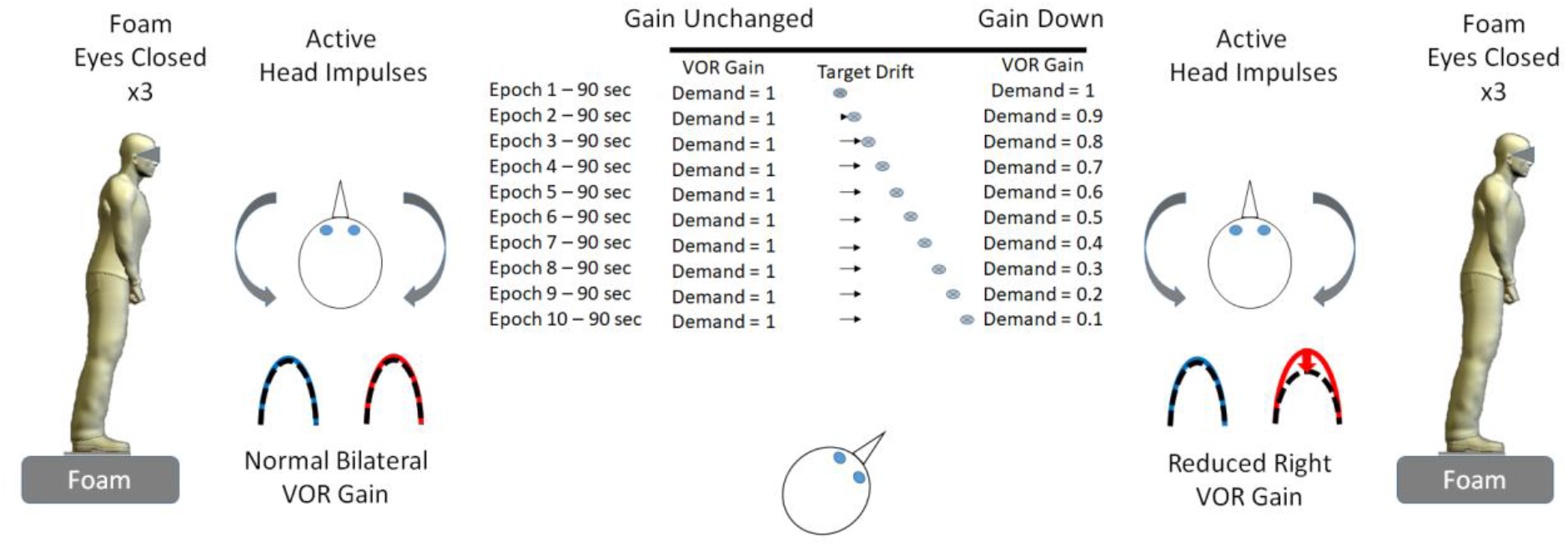
Experimental schematic visualizing the protocol. Sway area was measured three consecutive times (30 seconds each) with eyes closed on foam in a dark room. Subjects then sat and performed active head impulses in the dark for initial VOR gain measurement. Subjects then completed 15 minutes of unilateral horizontal gain down VOR adaptation. Subjects repeated active head impulses to characterize post-adaptation VOR gain. Sway area was measured three additional consecutive trials with eyes closed on foam in a dark room.

Active head impulses were performed such that subjects were seated in the dark 1.5 meters from the wall performed small (10-20 degree), fast (120-180 degrees/second) movements of their head in the yaw plane while fixating a point in space (projected laser dot). Subjects were taught to maintain approximately 15-20 degree chin tuck to align the horizontal SCC with the horizontal plane, but this was not verifiable throughout the experiment as the entire experiment was performed in a darkened room. Head impulses were performed until 15 impulses were accepted by the EyeSeeCam software.

The side of unilateral incremental VOR gain down adaptation was determined by coin-flip the day of the experiment. Following the initial set of seated active head impulses, the StableEyes system VOR gain demand was set to the subject’s measured VOR gain (rounded to the nearest 0.05) for right and left, and the side to be adapted was set to increment down by 0.1 every 90 seconds for 10 epochs.(Migliaccio and Schubert 2013) After the initial epoch, the laser target presented a decreasing demand of the VOR gain in the direction of VOR gain = 0.1 by drifting in the direction of head motion prior to extinguishing, see Figure 1. The VOR gain demand for the non-adapting side remained equal to the pre-testing value.

After completing the VOR adaptation protocol, subjects immediately performed active head impulses as described previously to measure post-adaptation VOR gain. Subjects then stood up onto the foam pad and closed their eyes (the room remained dark the entire time), and sway area was measured as described above for three repetitions.

### Data Analysis

VOR gain was calculated as the ratio of eye velocity to head velocity at 60ms after the onset of head impulses. Percentage change in VOR gain was determined and compared to 5% change as a threshold for meaningful change based on prior work.(Mahfuz et al. 2020) Individuals with less than 5% decrease in target side VOR gain were invited to repeat the experiment or were excluded from analysis. Sway area for the three pre-adaptation tests were averaged (Pre). Post-adaptation sway area was divided into Post1 (first 30 second test) and the average of the last two tests (Post2).

A repeated mixed model compared sway area across time (Pre, Post1, Post2) while accounting for within subject variance. Post-hoc pairwise comparisons were completed with Bonferroni correction for multiple comparisons. A t-test was used to compare percentage change in VOR gain to zero, with α = 0.05, and as a sensitivity analysis we repeated that t-test analysis comparing percentage change in VOR gain to - 5%. Exploratory paired t-tests were performed to compare the percentage VOR gain change for the adapted and non-adapted side in a subset (n=6) of subjects who performed seated active head impulses again after the balance testing was completed. This post-hoc exploratory analysis was conducted to see whether the VOR gain adaptation was lost after standing up.

## Results

Two female subjects were excluded from the analysis based on atypical VOR adaptation or atypical body sway. In one case, the subject experienced bilateral VOR gain adaptation with the non-adapting ear experiencing approximately 10% VOR gain reduction despite VOR gain demand did not increment. Although consistent with prior work showing contra-adapted gain changes (Migliaccio and Schubert 2013), this subject’s balance responses may reflect a dose dependent change in sway since bilateral vestibular hypofunction results in worse balance than unilateral vestibular hypofunction. The other case demonstrated a sway pattern inconsistent with standing, achieving a sway area of 52 degrees^2 suggesting the subject moved their feet while shifting weight, see outlier in Post3 in Figure 2.

**Figure 2.**
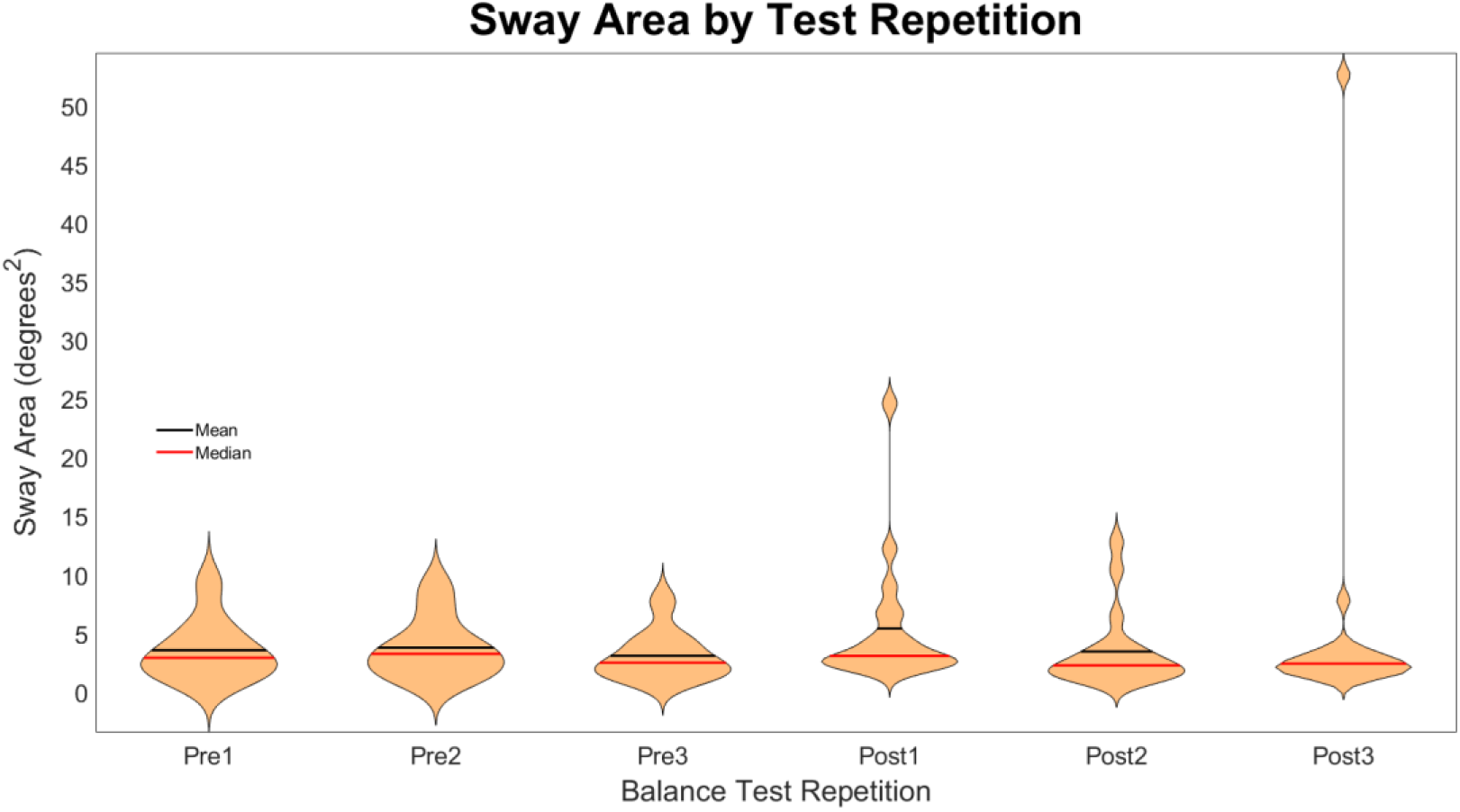
Violin plots for sway area for all subjects and all balance testing repetitions. Note the outlier in Post3 with sway area approximately 52 degrees^2, this subject was not included in statistical analyses. The black horizontal lines represent the group mean and the read lines represent the group median for each test repetition.

Average VOR gain measured from active head impulses before adaptation was 1.04 ± 0.05 on the left and 1.04 ± 0.07 on the right. Average VOR gain adaptation was - 10.9% ± 4.2%, individual responses are presented in Table 1. The reduction in VOR gain was significantly different from zero (t(1,15) = -10.4, p < 0.001, 95% CI [-13.1, - 8.7]). As a sensitivity test, we examined whether VOR reduction was significantly greater than 5%, a value previously identified as tolerance limit for the horizontal VOR (Mahfuz et al. 2020). VOR gain changed significantly more than 5%, (t(1,15) = -5.64, p < 0.001, 95% CI [-13.1, -8.7]).

**Table 1.** Individual subject VOR gain responses pre- and post-adaptation. Individuals marked with an * were excluded from analyses.

| Subject | Pre-adaptation<br>VOR Gain |  | Post-adaptation<br>VOR Gain |  | Percentage Change |  |
| --- | --- | --- | --- | --- | --- | --- |
|  | Left | Right | Left | Right | Left | Right |
| S124* | 0.92 | 0.98 | 0.81 | 0.88 | -11.9565 | -10.2041 |
| S1 | 1.04 | 1.17 | 0.89 | 1.1 | -14.4231 | -5.98291 |
| S128 | 1 | 1 | 1.05 | 0.94 | 5 | -6 |
| S103 | 0.98 | 1.02 | 0.94 | 0.87 | -4.08163 | -14.7059 |
| S122 | 1 | 1.02 | 0.98 | 0.92 | -2 | -9.80392 |
| S81 | 1.06 | 1.05 | 1.02 | 0.93 | -3.77358 | -11.4286 |
| S111 | 1.01 | 1.13 | 0.96 | 1.03 | -4.9505 | -8.84956 |
| S120 | 1.07 | 1.11 | 1.09 | 1.04 | 1.869159 | -6.30631 |
| S142 | 0.95 | 0.92 | 0.97 | 0.78 | 2.105263 | -15.2174 |
| S138 | 1.02 | 1.03 | 0.88 | 1 | -13.7255 | -2.91262 |
| S112 | 0.96 | 0.94 | 0.95 | 0.8 | -1.04167 | -14.8936 |
| S156 | 1.09 | 0.98 | 0.89 | 1.01 | -18.3486 | 3.061224 |
| S174 | 1.13 | 1.09 | 1.16 | 1.03 | 2.654867 | -5.50459 |
| S172 | 1.1 | 1.08 | 1.03 | 1.14 | -6.36364 | 5.555556 |
| S197 | 1.09 | 1.09 | 1.09 | 0.99 | 0 | -9.17431 |
| S202 | 1.07 | 0.99 | 0.92 | 1.02 | -14.0187 | 3.030303 |
| S171* | 1.17 | 1.14 | 1.03 | 1.21 | -11.9658 | 6.140351 |
| S164 | 1.03 | 1.04 | 0.97 | 1.07 | -5.82524 | 2.884615 |

Average pre-adaptation sway area was 3.2 ± 2.0 degrees^2, average Post1 sway area was 5.3 ± 5.8 degrees^2, and average Post2 sway area was 2.8 ± 2.2 degrees^2, see Figure 3. Planned post-hoc comparisons demonstrated an effect of time such that the Post1 sway area was significantly larger than Pre sway area (z = 2.61, p = 0.027, 95% CI [0.17, 4.12]) and Post2 sway area (z = -3.11, p = 0.006, 95% CI [-4.54, -0.59]).

**Figure 3.**
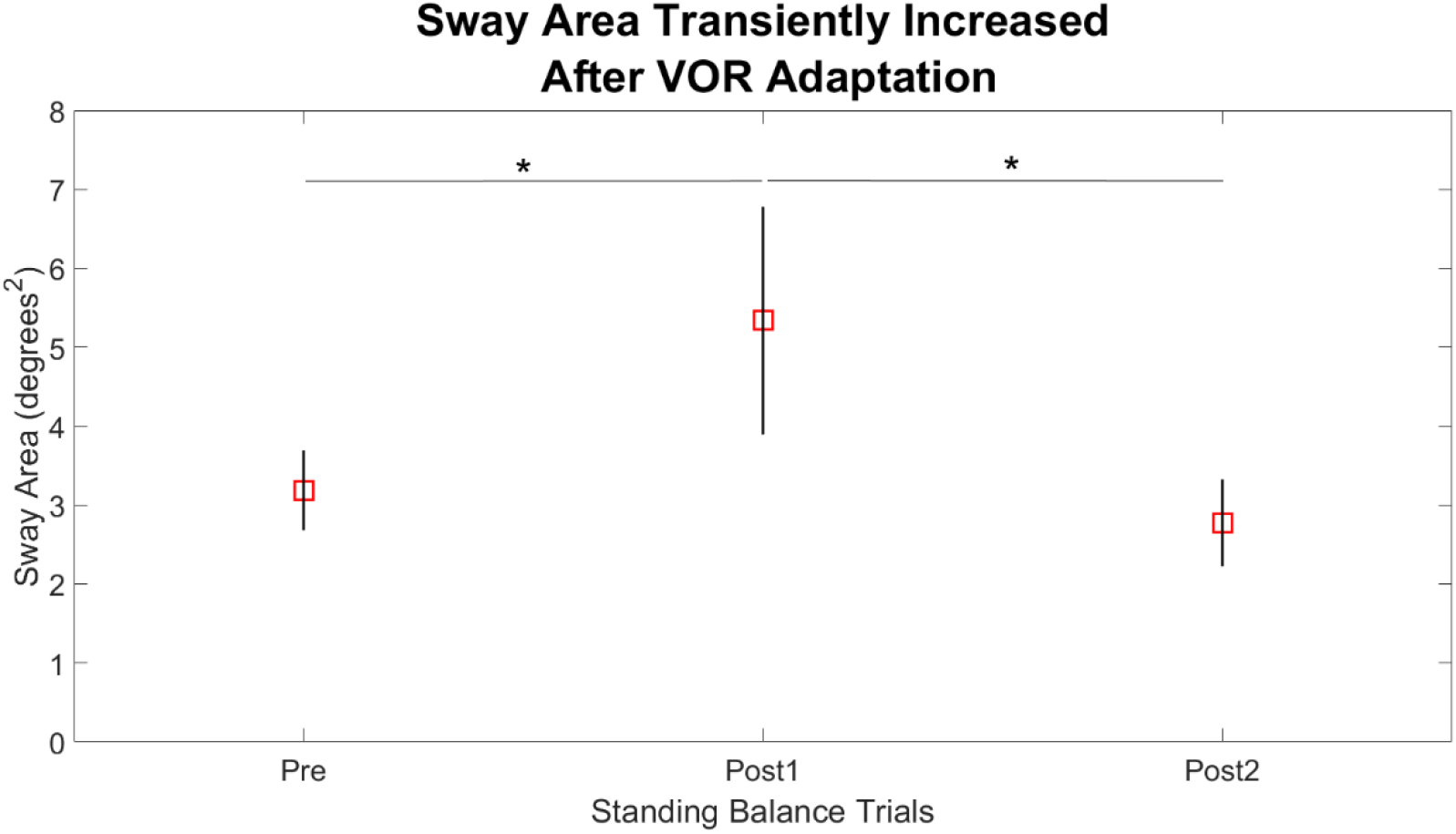
Average sway area before and after VOR adaptation. Error bars represent standard error of the mean. Note the significant increase in average sway area immediately after VOR gain down adaptation (Post1) that returns to baseline after the first post-test repetition (Post2), indicated by asterisk (p < 0.05).

In the subset analysis (n=6) examining whether VOR gain adaption was lost after standing, neither the adapted side (t(1,5) = 0.61, p = 0.57) nor the non-adapted side (t(1,5) = 1.47, p = 0.20) changed significantly. The average VOR gain percentage change on the adapted side before standing was -8.8% and after standing was -9.7%.

The average VOR gain percentage change on the non-adapted side before standing was 3.4% and after standing was 1.6%.

## Discussion

There are three primary results from the current study. First, we demonstrated an immediate worsening of postural control under a vestibular demanding postural task following unilateral VOR gain down adaptation. Second, we demonstrate a relatively fast normalization of postural sway consistent with sensory reweighting. Third, the sensory reweighting which corrected postural control occurred despite persistence of VOR adaptation.

Our current findings suggest in humans that semicircular canal projections appear to directly contribute to upright postural control. Vestibular signals diverge in the brainstem between pathways for the VOR and the vestibular only neurons involved in postural control (Cullen and Roy 2004; Cullen 2019). The results of this study indicate that the effects of the VOR adaptation paradigm are not restricted to the vestibulo-ocular pathways. Others have shown projections from semicircular canal pathways converge in the vestibular spinal system and influence canal plane specific head on body orientation and postural control (Suzuki and Cohen 1964; Shinoda et al. 2006).

However, projections from semicircular canals to the anti-gravity postural muscles may be less direct. Grillner and Hongo (1972) reported that otoliths but not semicircular canals projected to the extensor muscles in the trunk/lower limbs involved in postural control (Grillner and Hongo 1972). Both rotation and translation vestibular signals are represented in non-eye movement vestibular nuclei neurons (Newlands et al. 2018), and in cats ablation of semicircular canal function impaired postural responses above 0.1 Hz (Schor and Miller 1981). Both healthy adults and individuals with peripheral vestibular hypofunction demonstrate increased sway in the 0.1-1.0 Hz frequency band when standing on foam (Fujimoto et al. 2014). Thus, the observed sway increase may occur due to adapted semicircular canal signals combining with otolith signals prior to descending vestibulo-spinal tracts. Alternatively, the effects of angular VOR adaptation may directly modify otolith signaling. Future studies should examine whether rotational VOR adaptation leads to a change in otolith output.

Sensory weighting is a dynamic contextual process that facilitates upright stance under a variety of environmental and movement conditions (Assländer and Peterka 2016; Allison et al. 2018; Medendorp et al. 2018). Standing on foam is often described as a method to bias the reliable sensory input to the vestibular system (Cohen et al. 1993, 2019; Anson et al. 2017). Patients with vestibular hypofunction initially have difficulty standing on foam (Sprenger et al. 2017), but after rehabilitation (typically weeks to months) their ability to maintain balance on foam improves (Strupp et al. 1998; Giray et al. 2009). Our observed sway pattern, increased sway after a mild unilateral VOR gain down adaptation, is consistent with findings in unilateral vestibular hypofunction (Allum et al. 2017).

A possible explanation is reduced reliance on vestibular signals for postural control despite standing in the context of increased vestibular demand (foam with eyes closed). Reduced sway responses to vestibular signals when on foam with eyes closed occurred immediately after sub-concussive repetitive soccer heading which normalized 24 hours later (Hwang et al. 2016). The current results are consistent with a switch from normal integration of vestibular signals for postural control to a reduced reliance on of vestibular signals (immediate increased sway) followed by a change in the overall sensory weighting scheme to facilitate sway normalization. Future studies are needed to determine whether proprioceptive weighting increased, or whether converging semicircular canal and otolith signals was modified, or some combination explains the current results.

Interestingly, healthy young adults experience a small but significant sway increase across repeated tests of standing on foam, which corresponded to a shift from higher to lower frequency sway (Sozzi and Schieppati 2022). The rapid reduction (normalization) of sway area we observed suggests our subjects experienced a different process, more consistent sensory reweighting. Also in healthy subjects, Matsugi et al. (2017) attributed sway reduction on foam with eyes closed seen after performing 0.5Hz gaze stability exercises to a change in sensory strategy evidenced by a facilitation of the vestibulospinal reflex (Matsugi et al. 2017). Their results support our theory that sway normalization results from a change in sensory weighting; although in their study, the VOR gain was not adapted. Their results may alternatively represent enhancement in vestibular postural control due to priming, like the observed effect of prior eye movements enhancing the VOR (Das et al. 1999). Future studies should explore the dynamics of sensory weighting after VOR adaptation to determine the underlying mechanism. However, normalization of VOR gain response would also account for sway normalization in these healthy adults.

Support for sensory reweighting as the mechanism of sway reduction rather than normalization of the VOR is provided by additional testing in a subset of subjects. In a subset of subjects (n = 6) we retested active VOR gain after the post-adaptation balance testing was completed. All prior studies examining incremental VOR adaptation were performed in sitting and we conducted this sub-set analysis to determine whether the VOR adaptation was lost as a possible explanation for the sway normalization. In this subset, VOR gain on the adapted side remains reduced indicating the original adaptation did not wash out from the subjects standing and then sitting again. Others have also shown that gain down VOR adaptation in the pitch plane persisted in monkeys up to 7 days after adaptation (Schubert et al. 2008). Although our retention period was much shorter, our results suggest that the rapid sway normalization does not occur due to rapid recovery of adapted VOR. This is consistent with work in individuals with peripheral vestibular hypofunction who demonstrate improvement in postural control earlier than improvement in VOR responses (Allum et al. 2017). Future studies will need to examine whether similar results occur after gain up VOR adaptation. It is interesting to note that there is no nystagmus or dizziness following incremental VOR adaptation.

### Limitations

There are several limitations to the present results. Methodology limitations preclude definitive sensory reweighting based on sway responses to controlled sensory inputs, as reported in other studies (Peterka 2002; Hwang et al. 2014). However, the decline followed by improvement in postural performance while standing on foam is consistent with previous reports when vestibular sensory input is suddenly lost and later balance improves despite persistent vestibular hypofunction (Strupp et al. 1998; Allum 2012). Otolith function was not quantified in this healthy young adult cohort; although, no subjects had any neurological conditions and only one subject was over 60 years old (presbyvestibulopathy, Agrawal et al. 2019). Vertical VOR gain was not quantified and it is unknown whether there horizontal VOR gain adaptation results in any off-plane adaptation in pitch or roll. Small but significant changes in horizontal VOR gain were reported after pitch down gain change in healthy adults (Todd et al. 2022). Both roll-tilt and pitch are implicated in postural control (Wagner et al. 2021; Gabriel et al. 2022), and future studies should examine whether off-plane VOR adaptation occurs that may contribute to greater sway.

## Conclusion

These results indicate that adaptations targeting the lateral semicircular canal VOR pathways have broad postural effects. Rapid return of body sway to baseline levels despite persistent reduced VOR gain suggests dynamic sensory reweighting to resolve the transient but undesirable increased body sway. Normalization of body sway occurred without normalization of VOR gain suggesting the postural control system may be less tolerant to mild deviations from typical performance.

## Acknowledgement

We thank Professor Americo A. Migliaccio and Mr. Christopher J. Todd from the Balance and Vision Laboratory, Neuroscience Research Australia, for building the StableEyes devices and related software used in this study

